# Fast and alignment-free flavivirus classification from low-coverage genomes

**DOI:** 10.64898/2026.02.20.706982

**Authors:** Afsheen Shahid, Jens-Uwe Ulrich, Denise Kühnert

## Abstract

High genomic variability among viral species makes sequence classification highly dependent on multiple sequence alignment (MSA) methods, which are both computationally intensive and sensitive to data quality issues. To provide a more efficient and robust alternative, we developed DiCNN-UniK, a Dual-Input Convolutional Neural Network (DiCNN) utilizing unique ***k***-mer signatures and universal ***k***-mer libraries to generate novel and direct embeddings. Instead of relying on ***k***-mer frequency patterns, DiCNN-UniK directly leverages ***k***-mer embedding information, which provides a clear picture of local genomic context. This architecture is designed to handle full-length genomic sequences, overcoming the restrictive **512**-token limit common in many genomic foundation models. Trained on Flaviviruses, our model shows high sensitivity, robustness, and reliability, achieving an accuracy of **99%** on an independent test set. DiCNN-UniK is trained on full-genome data and is able to handle partial genomic sequences without preprocessing, maintaining high accuracy and precision with genomic coverage as low as **20%**. DiCNN-UniK currently stands as the best available model for the classification of flaviviruses, offering a sensitive, robust, and reliable solution for sequence analysis under real-world genomic coverage and data quality scenarios.

## 1 Introduction

The changing climate and globalization facilitate the occurrence and recurrence of infectious diseases and pandemic-like situations across the globe. According to the WHO, approximately 17% of all emerging infectious diseases worldwide are vector-borne [1]. Historically, flaviviruses are responsible for several large-scale global epidemics [2] and currently affect almost 400 million people worldwide every year [3], resulting in a substantial number of life-threatening cases [2] and fatalities [4]. Flaviviruses are single-stranded positive-sense RNA viruses with a length of 10.5 to 11.5kb (kilobases) and a structurally conserved genome that translates into a single polyprotein [5]. The clinical symptoms of widely circulating flavivirus infections are relatively non-specific [6], and misdiagnosis is a common problem [7]. Accurate identification of the disease-causing virus is crucial for disease management and outbreak control. In a globalized world, diseases and vectors are circulating regularly, and increasingly favorable climate conditions enable the establishment of disease vectors in new habitats where they could not survive before. With the rapidly evolving landscape of viral infections around the globe, genomic surveillance is becoming the backbone of effective pandemic management, and an early and accurate identification of emerging virus variants is an integral part of this [8, 9].

Multiple sequence alignment has been the foundation for genomic identification, classification tasks and comparative sequence analysis [10]. However, the complexity and computational costs of MSA-based approaches increases with an increasing number of samples and the length of the query sequences, with error propagation in progressive alignments [11]. Classical MSA-based bioinformatics tools also have limitations in providing accurate results when applied to real-world genomic data that is interspersed with ambiguous characters and incomplete information [12]

The genomic language, comprising of a defined alphabet, is similar to natural languages [13]. There are, however, no clear or predefined words or length of words in a genome. The concept of dividing the genomic sequences into substrings of defined lengths also referred to as ‘short-words’ was introduced by Claude Shannon in the form of “N-Grams” [14] which evolved into “*k*-mers” by the late 1990’s [15, 16]. The two prerequisites for developing a suitable vocabulary for genomic classification tasks are (i) the ideal length of the words/*k*-mers that can capture the full genomic pattern without losing important features (size too small) or causing heterogeneity (size too large) and (ii) the right balance between unique and common *k*-mers at the full genome level. Substrings of genomic sequences with a defined length of k have been utilized for pre-analysis in MSA tasks [17, 18] and have enabled addressing the challenges of conventional analytical methods in full genome analyses [18].

The utility of language foundation models based on natural language processing (NLP) for genomic downstream tasks, such as sequence classification and generation [19], has grown manifold over the last few decades. Recent years have seen the development of several unique strategies for different machine learning models, to define optimum ‘words’ and/or length of ‘words’ in a given corpus of genomic sequences. For example, the genomic foundation model DNABERT-1 [20] uses *k*-mers of different sizes (3-6), Nucleotide Transformer [21] uses only *k*-mers of size 6, GROVER [22] and DNABERT-2 [23] work with byte-pair encoding, and HyenaDNA [24] inputs are single nucleotides. DNABERT-1 [23] and Nucleotide Transformer [21] are foundation models that use *k*-mers as input to transformer-based models for genomic classification. The pre-existing versions can be trained on custom datasets and used as transfer models for different biological entities. However, they are technically constrained by the quadratic scaling of the self-attention mechanism, resulting in computational costs growing by *O*(*L*^2^) relative to the sequence length *L* [24, 25]. Many existing models, such as DNABERT-1 and Nucleotide Transformer, are pre-trained with a fixed context window length of typically 512 tokens. To train on the full-length Flavivirus genome (≈ 10,500–11,500 nucleotides) with these foundation models, the sequences need to be either truncated, which can cause significant loss of critical sequence information, or divided by utilizing a sliding window approach. Segmenting the genome into independent 512-token fragments on one side can disrupt the continuity of long-range gene features, and on the other side, will increase the architectural complexity required to aggregate local predictions of multiple sub-models into a cumulative classification result. Currently, there is no working system that deals with the genomic classification of circulating flaviviruses in Europe as a collective attempt. Our aim is to develop a robust classification model that can be trained on thousands of nucleotides/base pairs and can handle the inference of real-world data comprising incomplete and ambiguous genomic sequences. Considering *k*-mers in a nucleotide string comparable to small words in a text corpus, the linguistics power laws can be applied on sequencing data [26], as they apply to real-world languages [27], to find the ‘Hapax-Legomenon’ or ‘unique *k*-mers’. We propose that the combination of unique and common *k*-mers provides a fingerprint for virus classification at the species (and DENV serotype) level.

In this study, we build a novel classification model for flavivirus sequences using custom *k*-mers without multiple sequence alignments or pre-trained embeddings from existing genome foundation models. Our approach is based on a simple analytical solution for finding the optimal *k*-mer size for the selected viruses. By utilizing this statistical linguistics approach and applying the language corpus power-laws on the viral genome reference sequences we developed a genomic vocabulary for optimal *k*-mer sizes based on universal *k*-mer libraries. We used this vocabulary to develop a robust classification model based on a dual-input convolutional neural network architecture. The model is easy to train on thousands of base pairs and is effectively handling the inference of real-world data comprising incomplete and ambiguous genomic sequences.

## 2 Results

We developed the DiCNN-UniK pipeline (Fig. 1) to perform virus classification. Our dataset included ten flavivirus classes, comprising four Dengue serotypes and six viruses known to circulate in the European region, namely yellow fever virus, Zika virus, West Nile virus, Usutu virus, tick-borne encephalitis virus and Japanese encephalitis virus. The number of samples was highly imbalanced, both for the genomic sequence coverage of samples from the NCBI-Virus database (see Fig. 3 (a)), and for the virus classes in the training dataset (see Fig. 3 (b)).

**Fig. 1:**
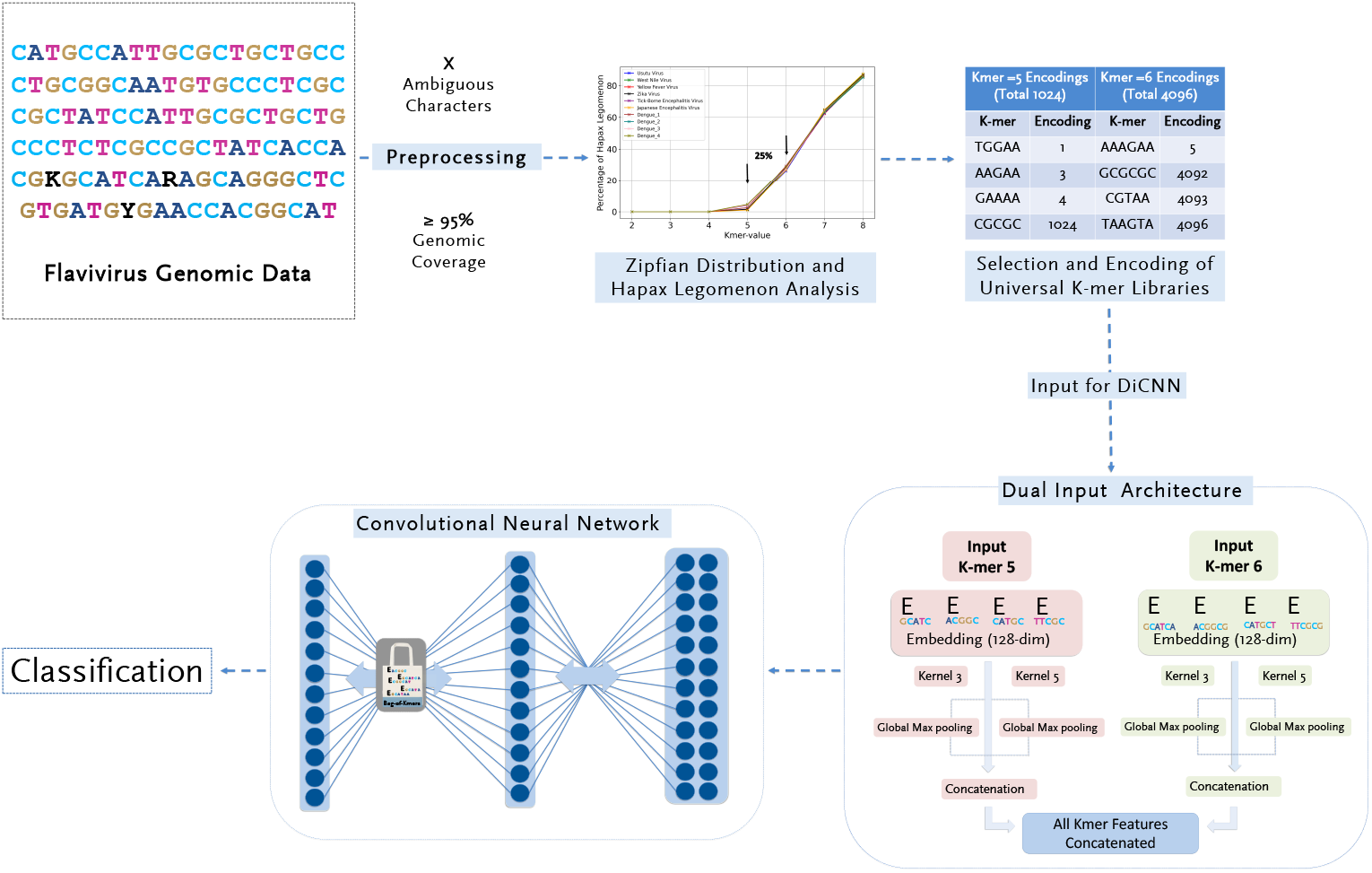
The DiCNN-UniK pipeline: **D**ual **i**nput **C**onvolutional **N**eural **N**etwork based on **Uni**versal *K*-mer libraries for multiclass flavivirus classification. *K*-mer sizes were optimized for capturing 25% unique signature via Zipfian Distribution and Hapax legomenon analysis.

### 2.1 Hapax Legomenon and selection of optimal *k*-mer sizes

We subjected the 10 selected flavivirus sequences (see Supplementary Table **??**) to Zipf’s law and calculated the percentage of hapax legomenon for *k*-mer sizes 2 to 8. As shown in Figure 2, the percentage of Hapax legomenon for a *k*-mer size of 5 is less than 10% and for a *k*-mer size of 6 is approximately 30% for all 10 flavivirus classes. A value lower than five results in a negligible amount of unique *k*-mers, and a value greater than six results in more than 50% of the *k*-mers being unique. For the selected flavivirus classes, we therefore chose a balance of 25% unique and 75% common *k*-mers, which occurs between sizes *k* = 5 and *k* = 6, and both were selected to generate *k*-mer libraries of corresponding sizes for input embeddings.

**Fig. 2:**
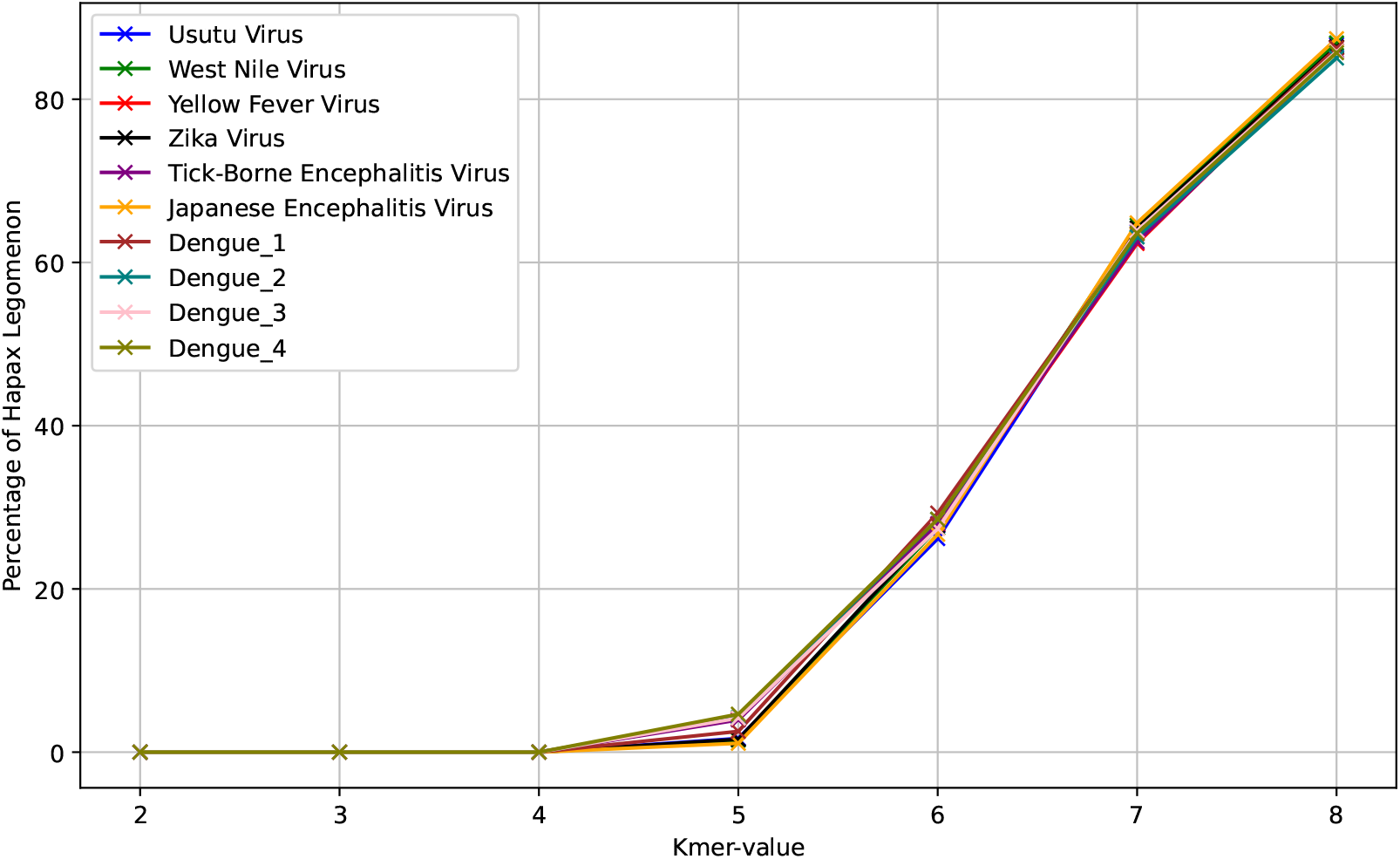
The percentage of Unique *k*-mers: ‘Hapax Legomenon’ with increasing *k*-mer size for 10 selected Flavivirus classes.

**Fig. 3:**
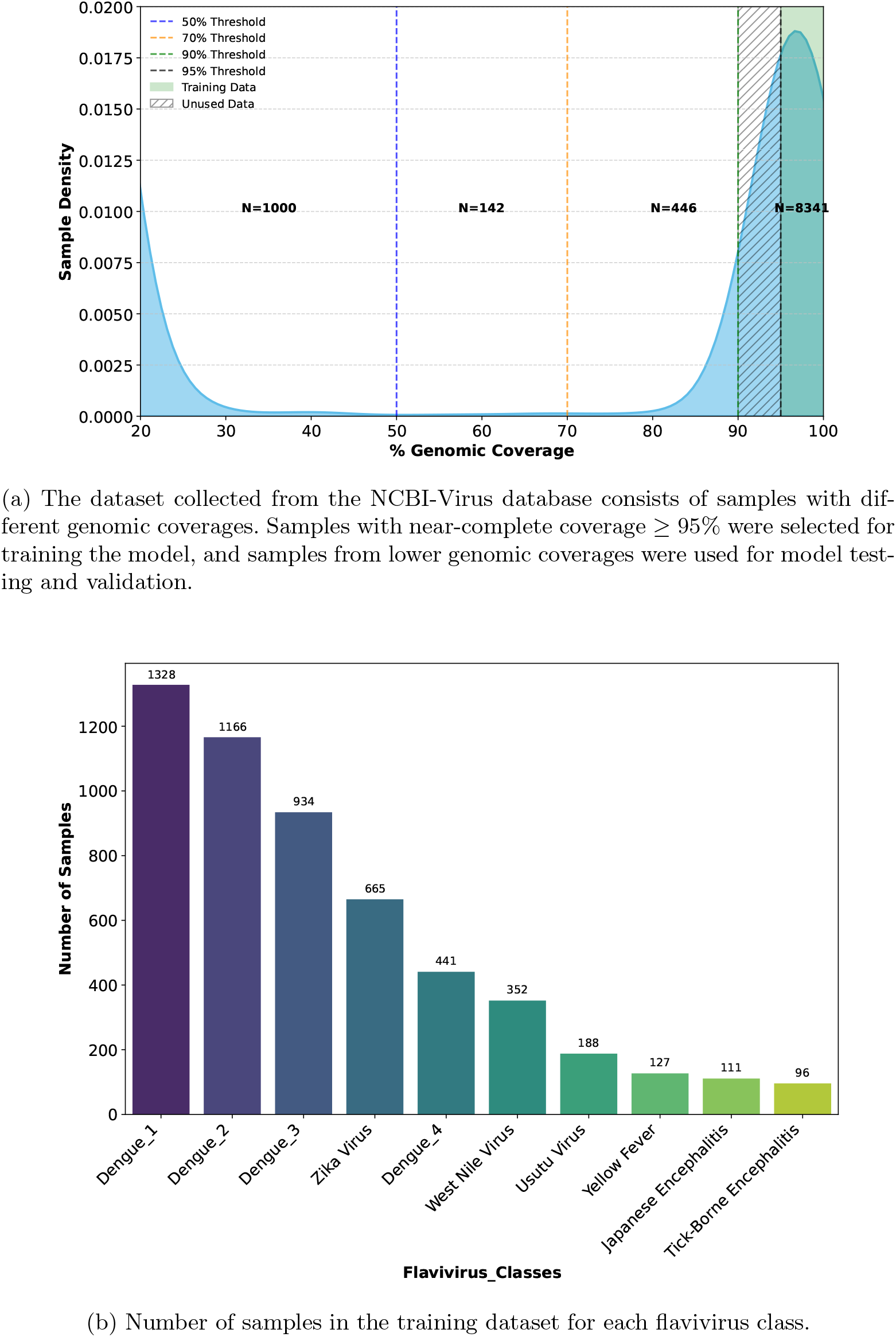
Data distribution for (a) Genomic Coverage of samples from complete NCBI-Virus dataset collected in May, 2025 (b) Virus Classes in the training dataset.

### 2.2 DiCNN-UniK: Training and Testing

The model was trained for 10 epochs on a local machine (see section 4.8) running for approximately 8 hours, using 6,672 clean samples (sequence composition of only ATGC) with a genome coverage of at least 95%, after the internal train-test split of 80-20. Following the initial training, the model’s performance was first evaluated on an independent test dataset of 1,669 samples. The model demonstrated strong predictive ability on test samples with 99% accuracy and an AUC value of 1.0. The corresponding confusion matrix confirms high predictive power, and the ROC curve shows a sharp ascent to the top left corner, demonstrating the model’s ability to maintain high sensitivity for all ten flavivirus classes (see Fig 4).

**Fig. 4:**
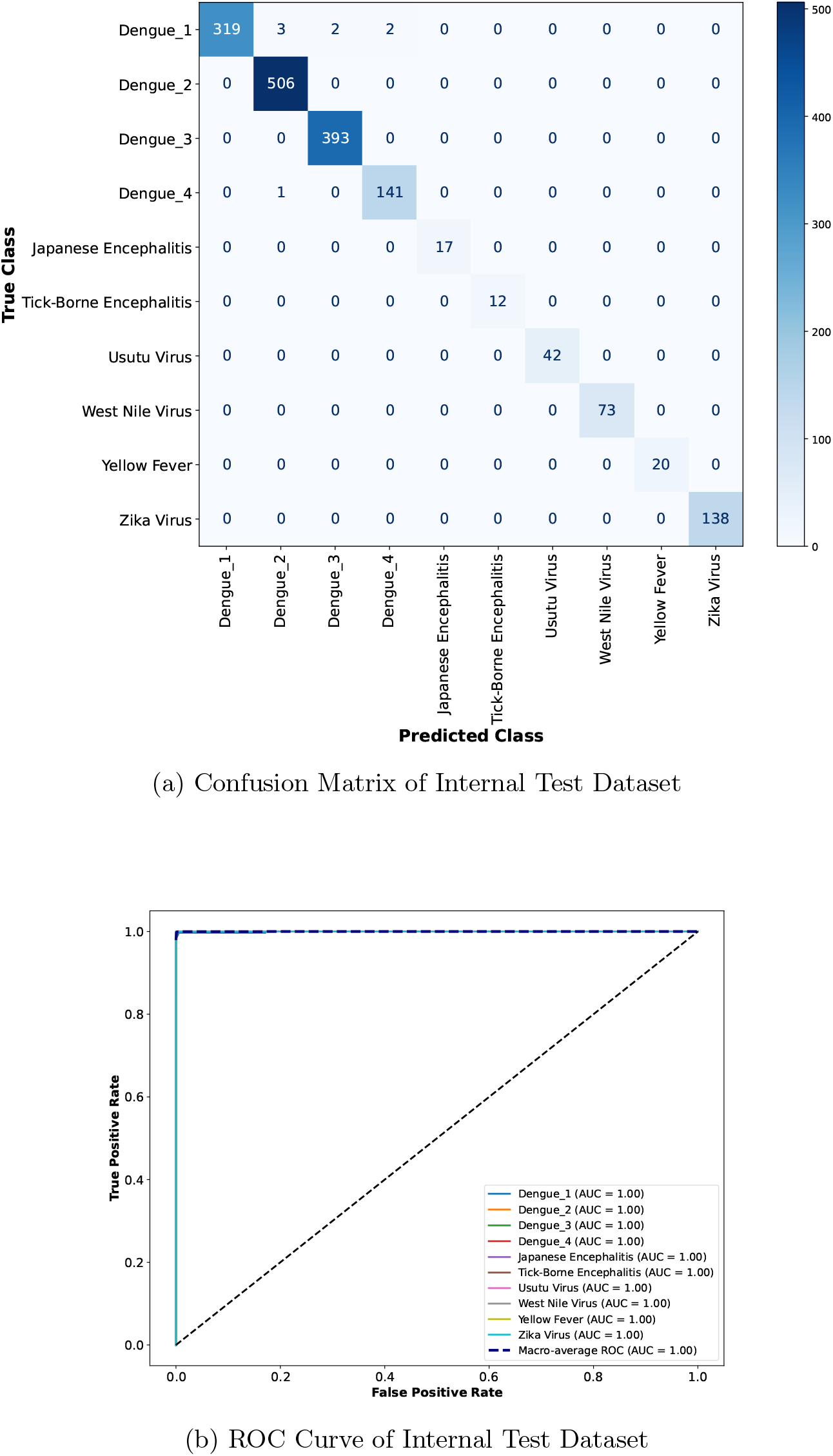
Performance of DiCNN-UniK shown through the (a) confusion matrix and (b) the ROC curve of the same internal test set.

### 2.3 External Validation with decreased genome coverage and unprocessed datasets

The model was externally validated on unprocessed samples (sequences containing IUPAC ambiguity codes for incomplete nucleic acid specification) with varied genomic coverages (20-70%, Fig. [3 (a)]). Although the model was trained on clean genomic data containing only A, C, G and T nucleotides, it performed the multiclass classification task without any errors when we tested the model on samples with as many as 9 different ambiguous characters (see Table 1). In fact, the model drops all *k*-mers that are not part of the universal *k*-mer library. Due to the one stride overlapping *k*-mer generation, only the ambiguous characters get dropped. Hence, the model efficiently cleans partial and full sequences without any pre-processing. Furthermore, we gradually and systematically decreased the genomic coverage of the validation datasets (see Table 1) to explore the sensitivity and robustness of our model when subjected to incomplete sequence data. There was no noticeable loss in accuracy and Matthews Correlation Coefficient (MCC) values when the genomic coverage reduced to 70%. A slight reduction was observed when genomic coverage was reduced to 50% and the model’s performance was stable for majority of flavivirus classes (Supplementary Figure **??**) even when the genome coverage was reduced to 20% (Supplementary Figure **??**).

**Table 1:** Validation datasets: qualitative and quantitative features.

| Dataset | Genomic Coverage (%) | Number of Samples | Unique Ambiguous Characters | Ambiguous Character Types (%) |
| --- | --- | --- | --- | --- |
| Test_set.1 | 70–90 | 446 | 9 | 9.46 |
| Test_set.2 | 50–70 | 142 | 6 | 7.09 |
| Test_set.3 | 20–50 | 1000 | 7 | 1.17 |

*The external validation test sets were curated from the NCBI database and divided into selected genomic coverages using post-filtering methods.*

### 2.4 Comparison of HyenaDNA Transfer Model and DiCNN-UniK

After external validation, we also compared the performance of our model with that of the most suitable foundation model. For this comparison, we selected HyenaDNA pretrained small–32k [24], the only available pretrained version of a genomic foundation model that could be trained for the sequence length of a full flavivirus genome of approximately 11,500 nucleotides. The HyenaDNA transfer model (HyenaDNA-TM) was trained for three epochs using the same training dataset (6,672 samples) and evaluated on the same internal test dataset (1,669 samples). This resulted in an accuracy of 99% and an AUC value of 1.0, making the training and internal testing performance comparable to DiCNN-UniK (Table 2).

**Table 2:** Training and validation performance comparison of HyenaDNA transfer model (HyenaDNA-TM) and DiCNN-UniK.

| Model | Training Time (m:s) | Number of Epochs | Trainable Params | Testing Accuracy | AUC Value | Inf. Time (ms) |
| --- | --- | --- | --- | --- | --- | --- |
| HyenaDNA-TM (Pre-trained) | 43:07 | 3 (Fine-Tuning) | 3,278,848 | 99% | 1.0 | 64.46 |
| Di-CNN UniK | 22:04 | 10 (Fine-Tuning) | 1,839,626 | 99% | 1.0 | 4.19 |

There is a clear difference in training duration for HyenaDNA-TM (3 epochs) versus DiCNN-UniK (10 epochs). HyenaDNA-TM rapidly reaches a higher performance because it is a pre-trained genomic foundation model that builds and develops the finetuning process with a robust internal representation of nucleotides. The model training performance shown in Supplementary Table **??** demonstrates a higher accuracy of HyenaDNA-TM even from epoch 1, achieving a training loss of 0.0285 and a validation loss of 0.0233, comparable to the performance metrics achieved by DiCNN-UniK at its 10th epoch (Supplementary Table **??**). Both models reach a similar steady-state performance. Further training of the 3.2M-parameters of HyenaDNA-TM beyond three epochs would have increased the risk of overfitting to the specific training distribution without providing any improvement in validation performance.

However, when we compared the performance of HyenaDNA-TM on external validation datasets with decreasing genome coverages and unprocessed sequences, the accuracy and MCC values were consistently below 50% for all three datasets (Table 3 and Fig. 5). The superior performance of DiCNN-UniK over the HyenaDNA transfer model across varying genome coverages for flavivirus classification can be attributed to the architectural focus of the former on small, specific genetic ‘fingerprints’ rather than the entire genome. HyenaDNA-TM utilizes long-range convolutions to capture global genomic patterns and the performance is very sensitive to sequence discontinuities and lower coverages. In comparison, our dual-input *k*-mer approach (*k* = 5, 6) and parallel convolutional kernels (*F* = 3, 5) allow the model to scan for short, highly conserved patterns even when the input sequence is incomplete or unprocessed. This results in a highly accurate identification of flaviviruses even from incomplete and low-quality data. The confusion matrices and ROC-AUC curve plots for all three validation test sets in Supplementary Figure **??, ??** and **??** demonstrate the superior performance of DiCNN-UniK over HyenaDNA-TM.

**Table 3:** Performance Comparison of HyenaDNA Transfer Model and DiCNN-UniK Across Three Validation Datasets.

| Test | HyenaDNA Transfer Model |  |  |  |  | DiCNN-UniK |  |  |  |  |
| --- | --- | --- | --- | --- | --- | --- | --- | --- | --- | --- |
|  | Acc. | F1 | Rec. | MCC | AUC | Acc. | F1 | Rec. | MCC | AUC |
| Test_set_1 | 0.13 | 0.08 | 0.13 | 0.08 | 0.76 | 1.00 | 1.00 | 1.00 | 1.00 | 1.00 |
| Test_set_2 | 0.26 | 0.13 | 0.02 | 0.02 | 0.75 | 0.97 | 0.97 | 0.97 | 0.97 | 1.00 |
| Test_set_3 | 0.41 | 0.24 | 0.007 | 0.007 | 0.57 | 0.98 | 0.98 | 0.98 | 0.98 | 0.99 |

*Note: Acc.: Accuracy; Rec.: Recall; MCC: Matthews Correlation Coefficient.*

**Fig. 5:**
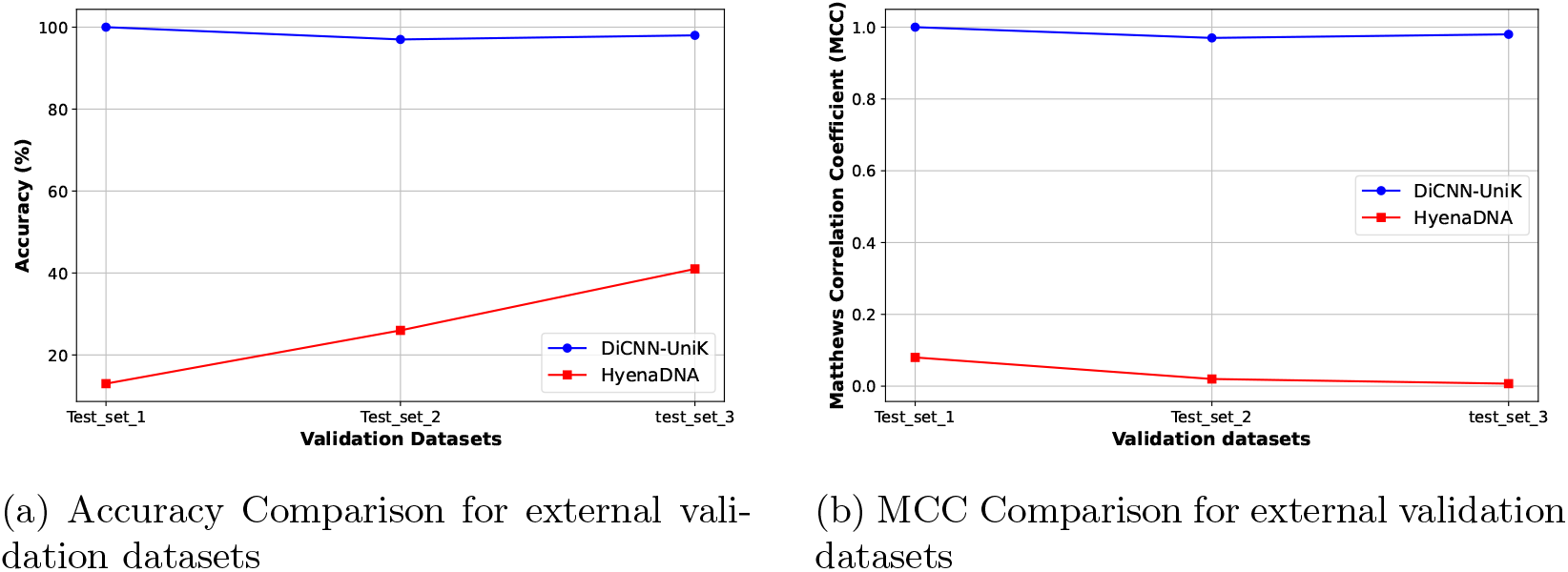
Performance Comparison between HyenaDNA and DiCNN-UniK (a) shows the accuracy, and (b) shows the MCC value comparison for all three external validation datsets.

In summary, DiCNN-UniK is highly sensitive, more robust and performed better than the HyenaDNA transfer model when subjected to unseen external validation datasets. It efficiently deals with unprocessed data containing ambiguous characters as well as sequences with low genomic coverage. Beside having the best classification performance, DiCNN-UniK needs substantially less time for training and inference while requiring only 56% of the trainable parameters of HyenaDNA-TM (Table 2), showing a great improvement also in computational performance.

## 3 Conclusion and Discussion

We developed a robust and sensitive multiclass classification model, based on universal *k*-mer libraries and unique *k*-mer patterns of flaviviruses, which handles reduced genome coverages without substantial loss in accuracy and precision. The architecture of DiCNN-UniK utilizes two important factors: the balanced utilization of unique and common *k*-mers and the ability of convolutional neural networks to extract important genomic features.

The high test and validation scores demonstrate the utility of our model, which combines *k*-mer representations and a deep convolutional neural network architecture to extract and identify signatures from unprocessed and incomplete genomic sequence data. To confirm the reproducibility and robustness of our model’s performance we tested it on three external validation datasets with varying degrees of genomic coverages without a noticeable loss in accuracy or MCC values. Instead of using frequency vectors, DiCNN-UniK directly encodes the *k*-mers. The CNN integrates these parts through non-linear activation functions between layers. This non-linearity allows the CNN to model complex, non-additive interactions between different *k*-mers. The convolutional architecture effectively translates the *k*-mer encoding distributions into classification features. The one-dimensional convolutional filters extract local dependencies and hierarchical relationships between *k*-mers, resulting in a context-aware and weighted representation of the most important *k*-mers as output of the CNN’s final dense layer.

Our results strongly support the hypothesis that unique *k*-mers carry a significant amount of information for fine-grained genomic classification. Common *k*-mers provide context at the family or genus level. However, their universal presence limits their utility for distinguishing between closely related strains or species within a single genus, such as different flaviviruses. In contrast, unique *k*-mers act as “signatures” that are selective, offering an informative pattern for classification. By defining a balancing point to obtain a sufficient number of unique and general *k*-mers, DiCNN-UniK captures the full spectrum of *k*-mer frequencies, allowing the CNN to learn the weights informed by unique and information-rich *k*-mers. This phenomenon closely follows Zipf’s Law [28], which describes the relationship between word frequencies and ranks in natural languages. In linguistic models, the most frequent words with the highest ranks, i.e. “the,” “and,” “a”, “is”, etc. are common but convey little meaning. The context is mainly defined by the less frequent nouns, verbs, and specific adjectives, which hold the lowest ranks. Similarly, the genomic sequence can also be interpreted as a text corpus. Common *k*-mers are the “stop words,” necessary for structure, but cannot define the full context. In contrast, unique *k*-mers are responsible for determining the identifying pattern. Our model extracts the highly specific “nouns” and “adjectives” by leveraging the unique *k*-mers of the viral genome, which are words that define its unique identity and function.

Our model represents a significant advance over both traditional and contemporary genomic classification methods. It shows a number of advantages over traditional methods and machine learning models, while surpassing the limitations of traditional alignment-based methods like BLAST [29] or phylogenetic analysis tools [30]. There are well-established alignment-free approaches such as KRAKEN [31] and *k*-mer frequency vector counting [32], which depend on *k*-mer frequency counts and do not look directly for unique sub-sequences or *k*-mers that distinguish closely related strains. While they are fast, they tend to lack the sensitivity required for fine-grained classification. The recently developed machine learning model Craft [33], which builds on alignment-based methods and BCGR feature representations, achieves high classification accuracy for Dengue subtypes at a fine-grained resolution, even under partial genomic coverages. However, DiCNN-UniK is more broadly applicable thanks to its multivirus classification at fine resolution, and does not require multiple sequence alignment. With that, our model enables a more efficient handling of incomplete sequences without relying on reference datasets. With existing deep learning and transformer based models, the token size limitation (generally 512 to 1024 tokens [20, 22, 23]) is a major hurdle when training transfer models on full length genomes for sensitive classification tasks. The DiCNN-UniK architecture utilizes a 1D-convolutional approach that scales linearly (*O*(*L*)), enabling the simultaneous processing of longer sequences without the memory overhead associated with large embedding vectors (e.g., 768 or 1024) and the quadratic scaling of the self-attention mechanism. While these large language models excel at broad sequence classification tasks, they are actually computationally intensive, comparatively slow, and critically drop in accuracy when dealing with low-quality, incomplete genomic data. Real-time clinical and surveillance systems typically have to deal with such data, which is an issue directly addressed with DiCNN-UniK through its unique and novel architecture finetuned to capture specific genomic “signatures”. Our model demonstrated superior performance compared to the large, generalized deep learning foundation model HyenaDNA [24]. The HyenaDNA transfer model required a considerably longer training time and a greater number of trainable parameters, yet achieved identical classification accuracy and AUC values with internal test datasets and performed very poorly when subjected to external validation datasets with reduced genome coverage. While foundation models like HyenaDNA excel at generalized sequence modeling, the high efficiency, reduced data quality requirements and low parameter count of our customized approach demonstrate that domain-specific, lightweight architectures can achieve better performance with significantly reduced computational requirements and faster inference time. The DiCNN-UniK model is a major improvement in classifying flaviviruses, showing both very high accuracy and great efficiency. The novel architecture uses both common and unique *k*-mers from the genome sequences. Our model can reliably identify different flaviviruses with an accuracy from 98% (with a minimum of 20% genomic coverage) to 100% (with *>* 70% genomic coverage), see Table 2, even with unprocessed genomic data (as many as 9 ambiguous character types, see Table 1). For the same validation datasets the accuracy of HyenaDNA-TM was between 13% to 41%, see Table 2. Additionally, our model is very efficient, requiring minimal computing power while delivering results extremely fast (in microseconds). This makes it a very practical tool for use in real-world systems like hospital labs and surveillance pipelines, allowing for quicker identification of diseases and faster responses to outbreaks. Based on the balanced representation of universal and unique *k*-mers, the architecture of DiCNN-UniK can be expanded as scalable foundation for future genomic analysis. DiCNN-UniK provides a blueprint for creating efficient, pathogen-specific classifiers that can outperform generalized foundation models in speed as well as sensitivity.

## 4 Methods

### 4.1 Data Curation

For each selected flavivirus (see Supplementary Table **??**), the accession numbers for the partial and full genomic sequences, along with the chosen metadata, were downloaded in comma separated values (CSV) format from the NCBI Virus database [34]. FASTA sequences corresponding to each accession number were recovered using Biopython and incorporated into the same files along with the associated metadata. For retrieving nucleotide sequences using accession numbers, the Biopython package (version 1.84) [35] coupled with the NCBI Entrez API [36] was used. Based on NCBI usage guidelines for pull request limits, accession numbers were fetched in batches of 50 using a custom wrapper function with a delay of 1 second between each batch request. The associated code and datasets are available on the github repository at https://github.com/AfsheenShahid/DiCNN-UniK/blob/main/Data.

### 4.2 Finding Duplicates

We used fuzzy string matching via the TheFuzz library [37] based on Levenshtein Distance [38] to identify and remove near-duplicate sequences between the training and external validation datasets to avoid data leakage. The de-duplication process was performed in two steps: First, a strict similarity threshold of 100% was applied, confirming that there were no exact duplicates between the datasets. Next, the threshold was relaxed to 95%, which is standard value in bioinformatics for reducing redundancy in genomic datasets [39], to identify potential near-duplicates. Sequences from the validation dataset that showed a similarity score above this threshold with any sequence from the training dataset were flagged as redundant. Once these high-similarity pairs were identified, any sequences in the validation set that were too similar to the training data were removed. This ensured that the validation dataset remained independent, with no overlap or contamination from the training sequences. The final validation datasets consisted of unique sequences only, preserving the integrity of the external validation process.

### 4.3 Zipf’s power law, Hapax Legomenon and *k*-mer Selection

*k*-mers are fixed-length short substrings of biological sequences. In bioinformatics research, *k*-mers are an important tool used for characterization and reconstruction of genomic sequences [17, 18]. *k*-mers in a given genomic sequence are comparable to words in a sample text and also follow the word frequency laws based on statistical linguistics [27]. Zipf’s Law [28, 40] is a simple mathematical distribution expressed by the equation

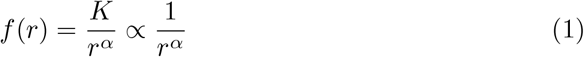

where *f* is the frequency of a word, *r* is its rank, and *K* and *α* are constants determined by the data. *K* is the constant of proportionality and a normalization factor. It is determined by the total number of words in the entire text corpus, in our case *k*-mers in the complete genome sequence. For instance, in a corpus with a total of *N k*-mers and *M* unique *k*-mers, *K* is defined such that the sum of all frequencies equals *N* [41] and calculated as:

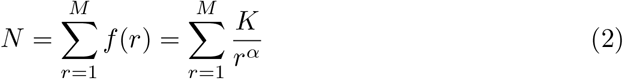

In turn, *K* is calculated as:

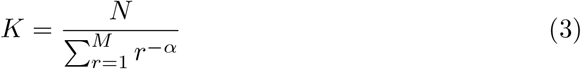

where the denominator is the generalized harmonic number of order *α*.

*α* is the Zipf’s exponent that determines the slope of the decreasing frequency or increasing rank of the *k*-mers depends on the constitution of the genomic sequence. Taking the natural logarithm of the Zipf’s equation yields:

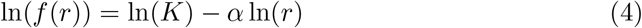

In this linear form, *α* represents the negative slope of the line. It is determined using the Least Squares Method or Maximum Likelihood Estimation (MLE) to fit the observed *k*-mer distribution [28]. Although Zipf’s power law is not a perfect fit for *k*-mers and genomic data [42], it can be used for finding the Hapax legomenon or the unique *k*-mers. Hapax legomenon are the unique words that appear only once in a given sample of text. The selection of an optimal *k*-mer size is crucial, as different sizes result in patterns with varying degrees of generalization and uniqueness within the genomic data. For example, a *k*-mer size of 3 represents universal codons and, in turn, is completely generalized for all living organisms. The selection of the size of the *k*-mers depends not only on the genomic makeup of the species of interest but also on the task, which in our case is a classification task to distinguish between different flaviviruses. This requires a specific balance between generalization and uniqueness. We utilized Zipf’s law and plotted the Zipfian distribution curve for *k*-mer sizes 2 to 8, and analyzed the content of the Hapax legomenon [43] for each size to find the optimal *k*-mer size for classifying flavivirus genomes.

### 4.4 *k*-mer Libraries and Encodings

We generated a universal library of *k*-mers that includes all possible *k*-mers for *k* = 5 and *k* = 6 over the nucleotide alphabet Σ = *{*A, T, G, C*}*, encoding the *k*-mers by using a simple integer encoding scheme. The total number of possible *k*-mers over the DNA alphabet Σ is given by 4^*k*^. For *k* = 5 and *k* = 6 the total number of 5- and 6-mers is

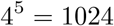

and

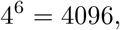

respectively. Table 4 represents a selection of these *k*-mers and their designated encodings. The full list of encodings is provided in the github repository at https://github.com/AfsheenShahid/DiCNN-UniK/blob/main/Kmer5_integer_encodings.csv, https://github.com/AfsheenShahid/DiCNN-UniK/blob/main/Kmer6_integer_encodings.csv.

**Table 4:** Selected *k*-mer Encodings.

| $k$-mer 5 Encoding (Total 1024) | | $k$-mer 6 Encoding (Total 4096) | |
| --- | --- | --- | --- |
| $k$-mer | Integer | $k$-mer | Integer |
| TGGAA | 1 | GGAAAA | 1 |
| GGAAA | 2 | ATGGAA | 2 |
| AAGAA | 3 | GGAAGA | 3 |
| GAAAA | 4 | TGGAAG | 4 |
| ATGGA | 5 | AAAGAA | 5 |
| ... |  |  |  |
| TCGGT | 1020 | GCGCGC | 4092 |
| CGTAA | 1021 | CGTACG | 4093 |
| TAAGT | 1022 | TAATCG | 4094 |
| GTCCG | 1023 | TAAGCG | 4095 |
| CGCGC | 1024 | TAAGTA | 4096 |

### 4.5 Embeddings

The integer tokens of the generated *k*-mers (sizes 5 and 6) were used as input for the TensorFlow Keras Embedding function in the neural network architecture. In this layer, we used a 2D tensor (sample size, input length) of positive integers as input to generate a 3D tensor (sample size, input length, output dim) with 128-dimensional feature vectors. The embeddings were generated independently for the *k* = 5 branch and the *k* = 6 branch. This process transforms the corresponding input tensors of lengths 11,560 and 11,559 into 3D tensors with a fixed depth of 128, providing 1,479,680 and 1,479,552 total latent features per sample, respectively. During model training, these 128 numbers for each *k*-mer were continuously adjusted via backpropagation for optimization.

### 4.6 Dual Input Convolutional Neural Network

The Convolutional Neural Network (CNN) architecture [44] was customized to incorporate the two *k*-mer branches as dual-input [45] see Supplementary Figure **??**.

The 128-dimensional vector embeddings are processed through a series of one-dimensional convolutional layers (Conv1D) with kernel sizes of *F* = 3 and *F* = 5 and both configured with 256 filters (total of 1024 filters across the entire model) to generate the corresponding high-dimensional feature maps, across both *k*-mer resolutions. All convolutional layers utilized a stride of 1 to ensure no loss of sequence information during sliding, and valid (no) padding was applied. These feature maps are then passed through Global Max Pooling operations to extract the most significant activation from each filter across the entire sequence. The pooled feature vectors from both convolutional kernels are concatenated to form a comprehensive representation for each *k*-mer size individually. These feature representations from both *k*-mer branches (5-mer and 6-mer) are later merged into a unified feature space, integrating sequence information at multiple *k*-mer size resolutions. Within the two *k*-mer branches, the kernel size 3 and kernel size 5 also look for the respective number of adjacent *k*-mers as a group. The combination of two *k*-mer branches and two kernels in each branch results in multiple resolution coverage by expanding the effective receptive field [46] through convolution. The effective receptive field (*L*_*eff*_) covered by a single filter output can be defined by the following equation:

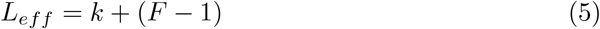

where *k* represents the input *k*-mer size and *F* represents the convolutional kernel size. Based on this relationship, the architecture achieves multi-resolution coverage as follows:

- Branch *k* = 5:
  – Kernel size *F* = 3 covers: 5 + (3 *−* 1) = 7 nucleotides.
  – Kernel size *F* = 5 covers: 5 + (5 *−* 1) = 9 nucleotides.
- Branch *k* = 6:
  – Kernel size *F* = 3 covers: 6 + (3 *−* 1) = 8 nucleotides.
  – Kernel size *F* = 5 covers: 6 + (5 *−* 1) = 10 nucleotides.

in this way, the combination of two *k*-mer branches and two kernels in each branch resulted in multiple resolution coverage of *k*-mer sizes 5, 6, 7, 8, 9, and 10. The resulting concatenated high-dimensional feature vector was then processed through a series of fully connected (dense) layers to capture nonlinear interactions among the extracted features. The first dense layer of 512 neurons converts the concatenated feature vectors into a more compact and discriminative representation, followed by a Rectified Linear Unit (ReLU) activation function [47] to introduce nonlinearity. A dropout layer with a 0.5 rate was also applied to prevent overfitting by randomly deactivating neurons during training, which enhanced generalization. The second dense layer, with 256 neurons, is followed by another dropout layer, which further reduces the feature dimensionality, maintains robustness and reduces dependency on specific neuron activations. The resulting output is passed to a softmax-activated output layer, where the number of neurons corresponds to the number of organism classes in the dataset. This layer converts the processed feature vector into a probability distribution across all potential classes.

### 4.7 HyenaDNA transfer Model (32k seq len pretrained version)

We trained a new transfer model based on the most suitable foundation model, HyenaDNA 32k seq len pretrained version from the LongSafari repository available at https://huggingface.co/LongSafari/hyenadna-small-32k-seqlen, using the same data DiCNN-UniK was trained on. The sequences were tokenized using HyenaDNA single-character tokenizer and the model was trained without changing the predefined default input parameters for the 32k seq len version with a maximum sequence length of 32,768 nucleotides, employing character-level tokenization across a 12-token vocabulary. Our HyenaDNA transfer model (HyenaDNA-TM) architecture is structured with 8 layers and a hidden dimension (*d*_*model*_) of 256, utilizing an expansion factor of 4 and a second-order Hyena operator. The source-code is available on the Github repository at https://github.com/AfsheenShahid/DiCNN-UniK/blob/main/hyenadna_tm_train_test_validate.py.

### 4.8 Hardware information

DiCNN-UniK was trained and validated on a local machine with 2 CPU cores and 12.67 GB of host RAM, running on an AMD64 architecture under Windows (Version 10.0.19045). The HyenaDNA transfer model was trained and validated on Google Colab utilizing NVIDIA Tesla T4 GPU with 15.36 GB of total memory under CUDA Version 12.4 and Driver Version 550.54.15. The GPU was confirmed to be operational and fully available to TensorFlow, which detected 1 physical device with 14.74 GB of usable memory. For comparison of training and inference speed, DiCNN-UniK training was repeated with the same hardware configurations as the HyenaDNA transfer model.

## Acknowledgements

The authors would like to thank Katharina Ladewig and Nils Körber for funding acquisition and Sandra Bütow for coordination support.

This work has been financially supported by the Germany Federal Ministry of Health (BMG) under grant No. 2523DAT400 (project “AI-assisted analysis and visualisation of pandemic situations” — AI-DAVis-PANDEMICS).

## Declarations

### Consent for publication

All authors have read and approved the final manuscript for publication.

### Data availability

The datasets generated and analyzed during the current study are available in the DiCNN-UniK repository at https://github.com/AfsheenShahid/DiCNN-UniK.

### Code availability

The custom DiCNN-UniK architecture and training scripts are open-source and can be accessed at https://github.com/AfsheenShahid/DiCNN-UniK.

## Author contribution

DK and AS conceptualized the study. AS designed the DiCNN-UniK architecture, performed the experiments and data analysis. All authors reviewed the architecture, analyses and results, and wrote the manuscript.

## Additional Information

### Supplementary Information

Available with this manuscript at [link to be provided by the publisher].

